# ARHGAP29 is required for keratinocyte proliferation and migration

**DOI:** 10.1101/2023.01.30.525978

**Authors:** Tanner Reeb, Lindsey Rhea, Emily Adelizzi, Bailey Garnica, Elliot Dunnwald, Martine Dunnwald

## Abstract

**BACKGROUND:** RhoA GTPase plays critical roles in actin cytoskeletal remodeling required for controlling a diverse range of cellular functions including cell proliferation, cell adhesions, migration and changes in cell shape. RhoA cycles between an active GTP-bound and an inactive GDP-bound form, a process that is regulated by guanine nucleotide exchange factors (GEFs), and GTPase-activating proteins (GAPs). ARHGAP29 is a GAP expressed in keratinocytes of the skin and is decreased in the absence of Interferon Regulator Factor 6, a critical regulator of cell proliferation and migration. However, the role for ARHGAP29 in keratinocyte biology is unknown.

**RESULTS:** Novel ARHGAP29 knockdown keratinocyte cell lines were generated using both CRISPR/Cas9 and shRNA technologies. Knockdown cells exhibited significant reduction of ARHGAP29 protein (50-80%) and displayed increased filamentous actin (stress fibers), phospho-myosin light chain (contractility), cell area and population doubling time. Furthermore, we found that ARHGAP29 knockdown keratinocytes displayed significant delays in scratch wound closure in both single cell and collective cell migration conditions. Particularly, our results show a reduction in path lengths, speed, directionality and persistence in keratinocytes with reduced ARHGAP29. The delay in scratch closure was rescued by both adding back ARHGAP29 or adding a ROCK inhibitor to ARHGAP29 knockdown cells.

**CONCLUSIONS:** These data demonstrate that ARHGAP29 is required for keratinocyte morphology, proliferation and migration mediated through the RhoA pathway.

## Introduction

The epidermis of the skin is mainly composed of epithelial cells (also known as keratinocytes) which undergo proliferation, migration and differentiation to maintain skin homeostasis, required for its essential barrier function.^1^ Particularly, keratinocyte migration is a highly dynamic process that allows a cell to respond to extracellular stimuli and rearrange its cytoskeleton.^2^ Many of the cellular behaviors required for migration are regulated by Rho family GTPases, including Rac, Cdc42, and Rho.^3^ These enzymes constantly shuttle between an active GTP-bound form, and an inactive GDP-bound form.^4^ Two classes of enzymes promote these changes: the Guanine Nucleotide Exchange Factor (GEFs) promote the GTP-bound form, and are considered activators of the GTPases, while the RhoGTPase Activating Protein (GAPs) promote the GTPase activity, and therefore are inactivating the GTPase. Cell migration is classically viewed with Rac promoting lamellipodia extension, Cdc42 promoting filopodial protrusion, and RhoA regulating actomyosin contraction and focal adhesions.^5^ While many more layers of complexity exist beyond this model, disrupting any of these key RhoGTPases can lead to extensive defects in cell morphology and migration.

We previously demonstrated that Interferon Regulatory Factor 6 (IRF6) regulates keratinocyte migration via the RhoA pathway.^6^ Particularly, we showed that *Irf6*-deficient murine keratinocytes exhibited increased active RhoA and contained more stress fibers by phalloidin staining.^6^ Treatment with a ROCK inhibitor, a downstream effector of RhoA, rescued the stress fiber and migration defects observed in these cells, indicating that the defects are Rho-dependent.^6^ In an attempt to determine the mechanism by which IRF6 regulates RhoA, analysis of microarray data on IRF6-deficient murine keratinocytes determined the most significantly differentially regulated RhoGTPase in IRF6-deficient skin was ARHGAP29. In addition to being one of the most highly expressed RhoGAP in the skin,^7^ *Arhgap29* is downregulated 1.5-fold in *Irf6*-deficient murine skin.^8^ Furthermore, *Irf6*-deficient keratinocytes displayed a reduction in ARHGAP29 by immunofluorescence and Western blot.^6,9^ These data demonstrate that ARHGAP29 is a downstream effector of IRF6, and could play a role in keratinocyte migration.

ARHGAP29 is a RhoGAP which preferentially regulates RhoA.^10^ It is required for embryonic viability, as homogygous loss of function murine embryos do not survive past mid-gestation, potentially due to a failure of chorioallantoic fusion.^11,12^ Consequently, not much is known about this enzyme. The majority of the studies evaluating the function of ARHGAP29 have been done in endothelial cells. In these cells, it reduces the activity of RhoA at the cell junctions to promote their remodeling required for tubulogenesis.^11,13–15^ ARHGAP29 is also upregulated in some cancer cells,^16–19^ and mediates the Yap-induced actin cytoskeleton remodeling during metastasis.^20^ A larger body of work has been done in relation to facial morphogenesis. In human, mutations in the *ARHGAP29* gene are associated with, and causative of, orofacial cleft.^9^,^21–27^ Consistent with this function, ARHGAP29 is detected in embryonic facial prominences, and mice heterozygous for *Arhgap29* exhibit orofacial defects.^12^ Given that cell proliferation and migration are required for palatal development, many genes associated with orofacial clefting are key regulators of proliferation and migration.^28^ However, despite all the data suggesting a role for ARHGAP29 in keratinocyte migration, its role in these cells remains untested.

In this study, we utilize CRISPR/Cas9 and small-hairpin RNA (shRNA) technologies targeted to *ARHGAP29* in a normal human keratinocyte cell line to determine the role of ARHGAP29 in keratinocyte biology. Keratinocytes with decreased levels of ARHGAP29 display altered morphology including an increase in cell area and an altered actin cytoskeleton as well as delayed population doubling time. Furthermore, we demonstrate that reducing ARHGAP29 protein levels delays keratinocyte migration following an *in vitro* scratch wound, demonstrating that ARHGAP29 is required for keratinocyte motility.

## Results

### Validation of Arhgap29 mutant keratinocyte cell lines

We took two complementary approaches to reduce the level of ARHGAP29 in keratinocytes. In the first approach, we transduced a keratinocyte cell line (Normal human skin keratinocytes, NHSK)^29^ with a lentivirus expressing CRISPR/Cas9 and either scrambled non-targeting gRNAs or gRNAs targeting exon 2 of *ARHGAP29*. Alleles were isolated by cloning exon 2 of *ARHGAP29*. Gene editing was confirmed following sequencing of this genomic region. The edited lines (CRISPR#1 and CRISPR#2) displayed deletions causing frameshift mutations (Supplemental Figure 1A). The frameshift mutations were predicted to cause premature truncation mutations within exon 2 leading to nonsense-mediated decay (Supplemental Figure 1B). In the second approach, we transduced NHSKs with lentiviruses containing small hairpin (sh) RNAs targeted to 3 different locations within the *ARHGAP29* gene (sh#1-3) as well as a non-targeting scrambled shRNA (control, shSc). One of the target is the 3’ untranslated region of the *ARHGAP29* mRNA (sh#3), making this cell line amenable to add-back the full length *ARHGAP29* cDNA for rescue purposes. Of the three shRNA lines, only sh#2 and sh#3 cells survived and grew; sh#1 cells were unable to proliferate and slowly died post-selection.

In order to validate the efficiency of our approaches, we performed Western blot analysis of protein extracts from these cell lines. Our results show that levels of ARHGAP29 protein were significantly decreased in edited keratinocytes compared to scrambled controls (range from 50 to 80 percent reduction, Figure 1A,B), yet some residual ARHGAP29 protein was detected in all cell lines. Add-back of ARHGAP29-GFP (+A29) in sh#3 keratinocytes was successful as demonstrated by the presence of a higher molecular weight protein due to the size of the GFP tag (Figure 1A).

**Figure 1:**
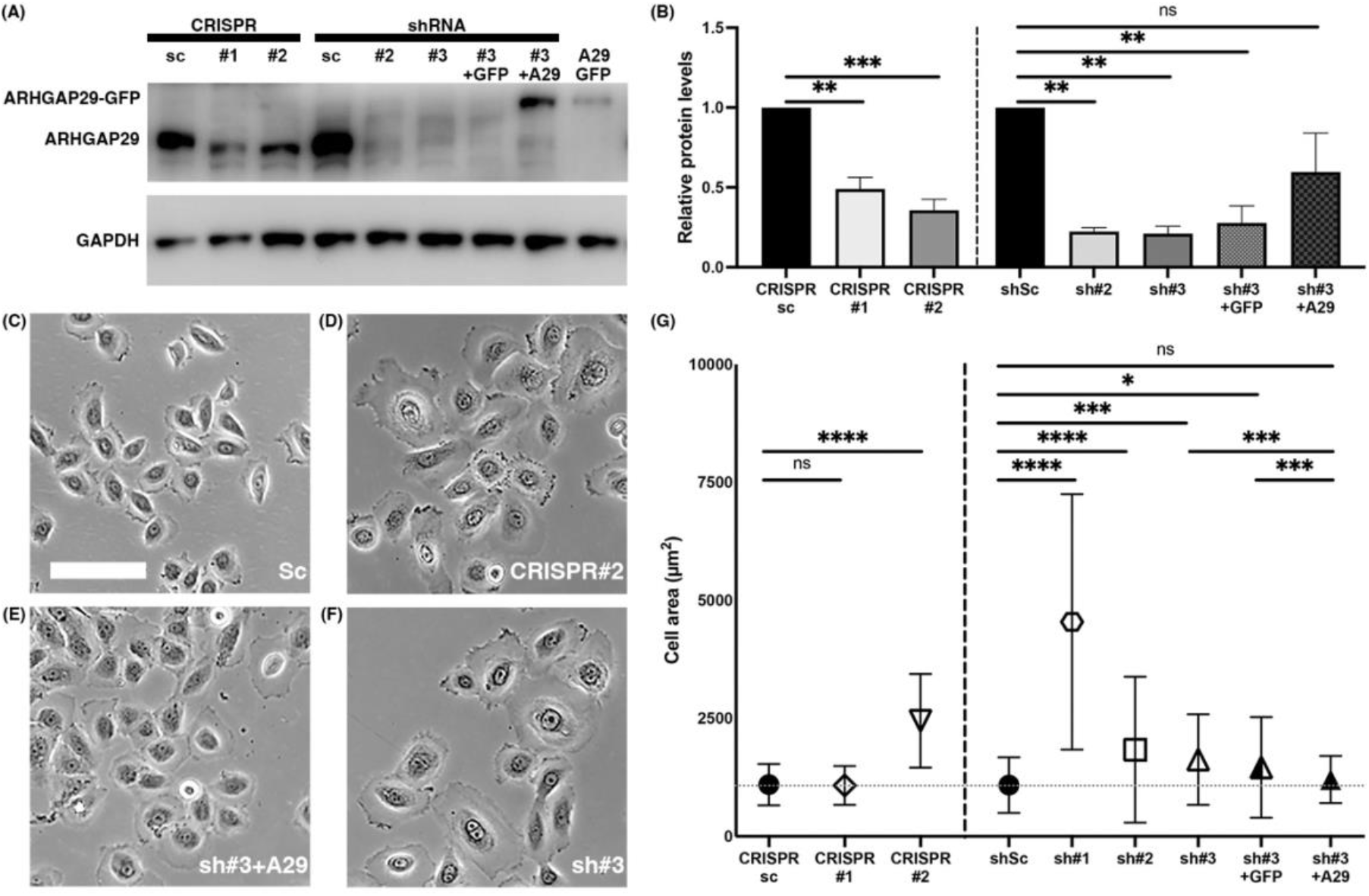
Characterization of Arhgap29 mutant keratinocytes. A, Western blot for ARHGAP29 and GAPDH (one representative of three experiments) of keratinocytes transduced with CRISPR scrambled (sc), CRISPR ARHGAP29 (#1, #2), shRNA scramble, shRNA ARHGAP29 (#2, #3), and HEK293T cells transduced with ARHGAP29-GFP (positive control). B, Quantification of ARHGAP29 protein levels. Values are the means (N = 3) ± SEM, ** *P* < 0.01 and *** *P* < 0.001 following ordinary one-way ANOVA test with Tukey’s multiple comparisons *post-hoc* test (only relevant comparisons are shown). C-F, Representative phase contrast micrographs of scramble (C), CRISPR knockdown (D), shRNA knockdown (F) and shRNA knockdown keratinocytes transduced with ARHGAP29-GFP (E), all grown in KSFM. Scale bar = 100 μm. (G) Quantification of keratinocyte area under the conditions shown in C-F. Values are the means ± SD, * *P* < 0.05, *** *P* < 0.001 and **** *P* < 0.0001 following Kruskal-Wallis test with Dunn’s multiple comparisons post-hoc test (only relevant comparisons shown). Red dotted line indicates the mean area of scrambled control keratinocytes. N = 200-400 cells per group. ns = non significant.

### Loss of ARHGAP29 increases keratinocyte size

ARHGAP29 is an inhibitor of RhoA, a major modulator of the actin cytoskeleton regulating cell shape.^10,30^ Therefore, we asked whether ARHGAP29 levels affect keratinocyte morphology. As shown in Figure 1C, control keratinocytes were refringent and exhibited smooth contours. However, cells with reduced levels of ARHGAP29 appeared less refringent, flatter, and with irregular contours compared to controls (Figure 1D,F). ARHGAP29 knockdown keratinocytes also appeared larger in size compared to controls (Figure 1D,F). Quantification of the area of the keratinocytes at 30-50% confluency showed a significant increase in cell area in ARHGAP29 knockdown keratinocytes compared to controls (Figure 1G). Interestingly, the largest increase in cell area was observed in the shRNA cells which were unable to proliferate *in vitro* (sh#1) and thus be studied further. The reintroduction of ARHGAP29 in sh#3 keratinocytes reduced the area of the cells to within the size of controls, demonstrating that this phenotype is specific to a reduction in ARHGAP29 protein levels (Figure 1E).

### ARHGAP29 levels affect downstream RhoA effectors

Cell shape and size are associated with increased active RhoA, which in turn increases actin polymerization (filamentous actin, or F-actin) and phosphorylation of myosin regulatory light chain (pMLC).^5,31^ In order to determine if ARHGAP29 levels affect these RhoA downstream effectors, we immunostained a subset of the knockdown keratinocyte cell lines for phalloidin, vinculin and phospho-myosin regulatory light chain (pMLC). Our results show that ARHGAP29 knockdown cells exhibited increased filamentous (F)-actin (as evidenced by phalloidin staining, Figure 1A-D) and pMLC (Figure 1C,D) compared to controls, suggesting that these cells may be more contractile. Collectively, these data demonstrate that reducing the levels of ARHGAP29 in keratinocytes promotes changes in the F-actin cytoskeleton.

### ARHGAP29 is required for keratinocyte proliferation

Studies have shown that small GTPases play a role in cancer and metastasis, including a potential role for ARHGAP29 in mediating Yap-induced actin cytoskeleton remodeling during metastasis.^20^ In fact, Rho GTPases and their activators (GEFs) and inactivators (GAPs) are known to directly regulate the cell cycle and cell proliferation.^32–34^ Therefore, we hypothesized that ARHGAP29 may regulate keratinocyte proliferation. To test this hypothesis, we calculated the population doubling time for each cell line. This assay estimates the time required for cells to double in number. Our results showed a significant increase in population doubling time in all ARHGAP29 knockdown cells compared to controls indicating a defect in proliferation (Figure 2A).

**Figure 2:**
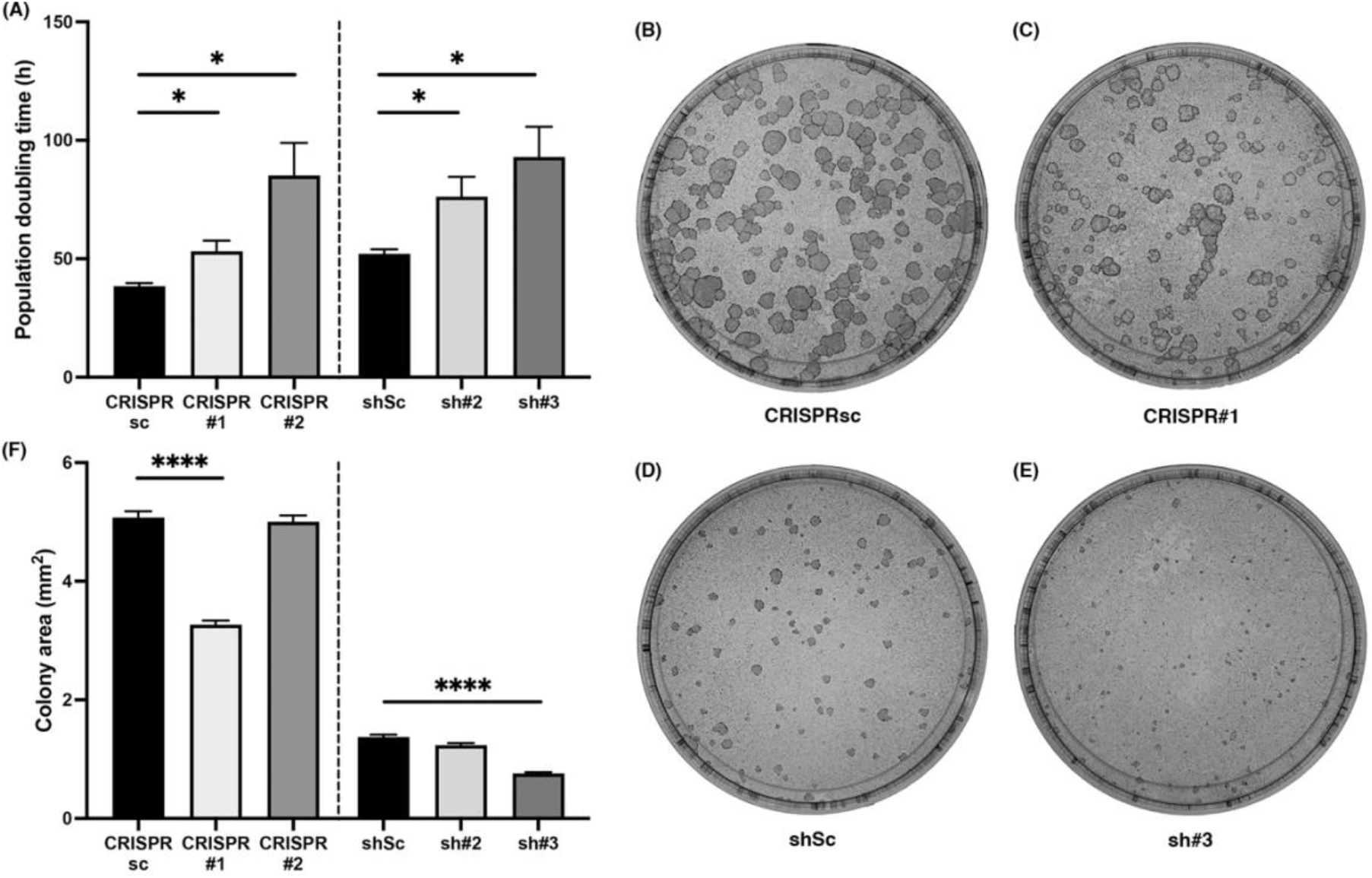
Arhgap29 is required for keratinocyte proliferation. A, Quantification of population doubling time. Values are the means ± SEM, * *P* < 0.05 following Brown-Forsythe test and Welch ANOVA test with Dunnett’s T3 multiple comparisons *post-hoc* test. N = 6-8 per group. B-E, Representative images of colony forming efficiency dishes with CRISPR scramble (sc, B), CRISPR ARHGAP29 (#1, C), shRNA scramble (sc, D) and shRNA ARHGAP29 (#3, E) keratinocytes. F, Quantification of colony area for all cell lines (including those represented in B-E). Values are the means ± SEM, **** *P* < 0.0001 following Kruskal-Wallis test with Dunn’s multiple comparisons *post-hoc* test (only comparisons to scrambled are shown).

To further investigate the proliferation defect observed in ARHGAP29 knockdown keratinocytes, we performed colony forming efficiency assays (CFE).^35^ First, we counted all colonies formed in the dish 2 weeks after plating. Our results showed no difference in the number of colonies between cell lines (data not shown) indicating all lines had the same potential for forming a colony. However, we observed smaller colonies in cell lines with reduced levels of ARHGAP29 (Figure 2B-F). Differences in colony size could be influenced by the number of cells in a colony or by the size of the cells in that colony. By measuring the area of all counted colonies, we found that only a subset of ARHGAP29 knockdown cell lines had significantly smaller colonies (Figure 2F). However, those that did not have significantly smaller colonies did have significantly larger cell size (Figure 1G). While the ARHGAP29 knockdown cell lines appear to have the same proliferative potential compared to controls, the decrease in colony area combined with the increase in cell size is consistent with the observed delay in population doubling time and supports a role for ARHGAP29 in proliferation.

### ARHGAP29 is required for single cell keratinocyte migration

RhoA is a key regulator of cellular migration.^36^ Previous work in fibroblasts showed that reduction in ARHGAP29 leads to impaired cell migration and scratch closure,^37^ however its role in regulating keratinocyte migration has not yet been explored. To directly test the hypothesis that reduction of ARHGAP29 impairs keratinocyte migration, we grew keratinocytes to confluency in low calcium medium, a condition in which calcium dependent cell-cell junctions are not fully engaged. Under these conditions, following *in vitro* scratch, keratinocytes moved as single cells to close the gap. Control keratinocytes closed the scratch within 24 hours following scratching (Figure 4A,C,E), however CRISPR ARHGAP29 keratinocytes displayed a delay in scratch closure (Figure 4B,D,E). These results demonstrate ARHGAP29 regulates keratinocyte migration.

**Figure 3:**
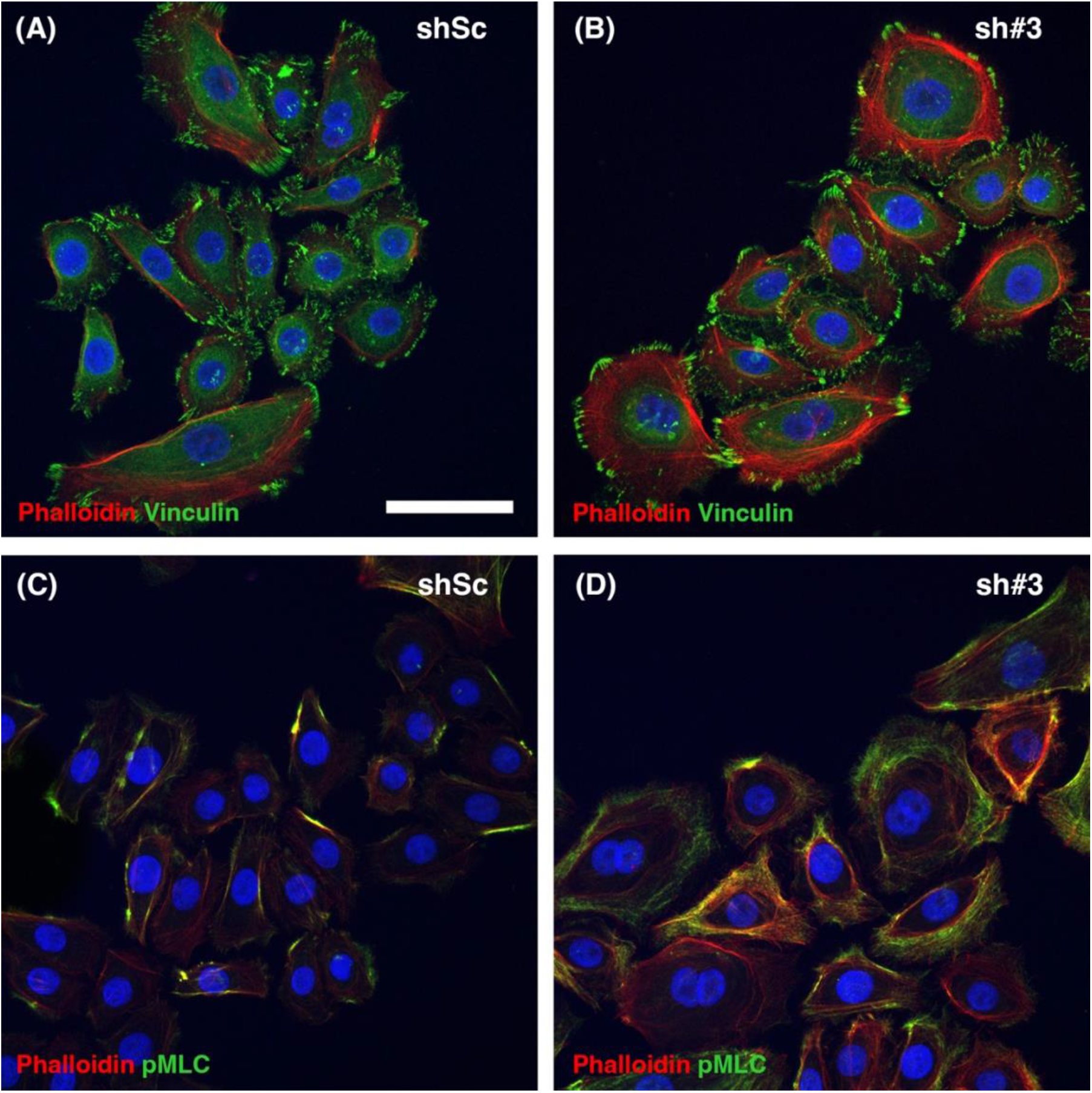
ARHGAP29 knockdown cell lines display increased stress fibers and phospho-myosin regulatory light chain (pMLC). Immunofluorescent staining for phalloidin (A-D, red), Vinculin (A,B, green) and pMLC (C,D, green) of shRNA scramble (shSc) keratinocytes (A,C) and shRNA ARHGAP29 (sh#3) keratinocytes (B,D). Nuclear DNA is stained with Hoechst (blue). Scale bar = 50 μm.

**Figure 4:**
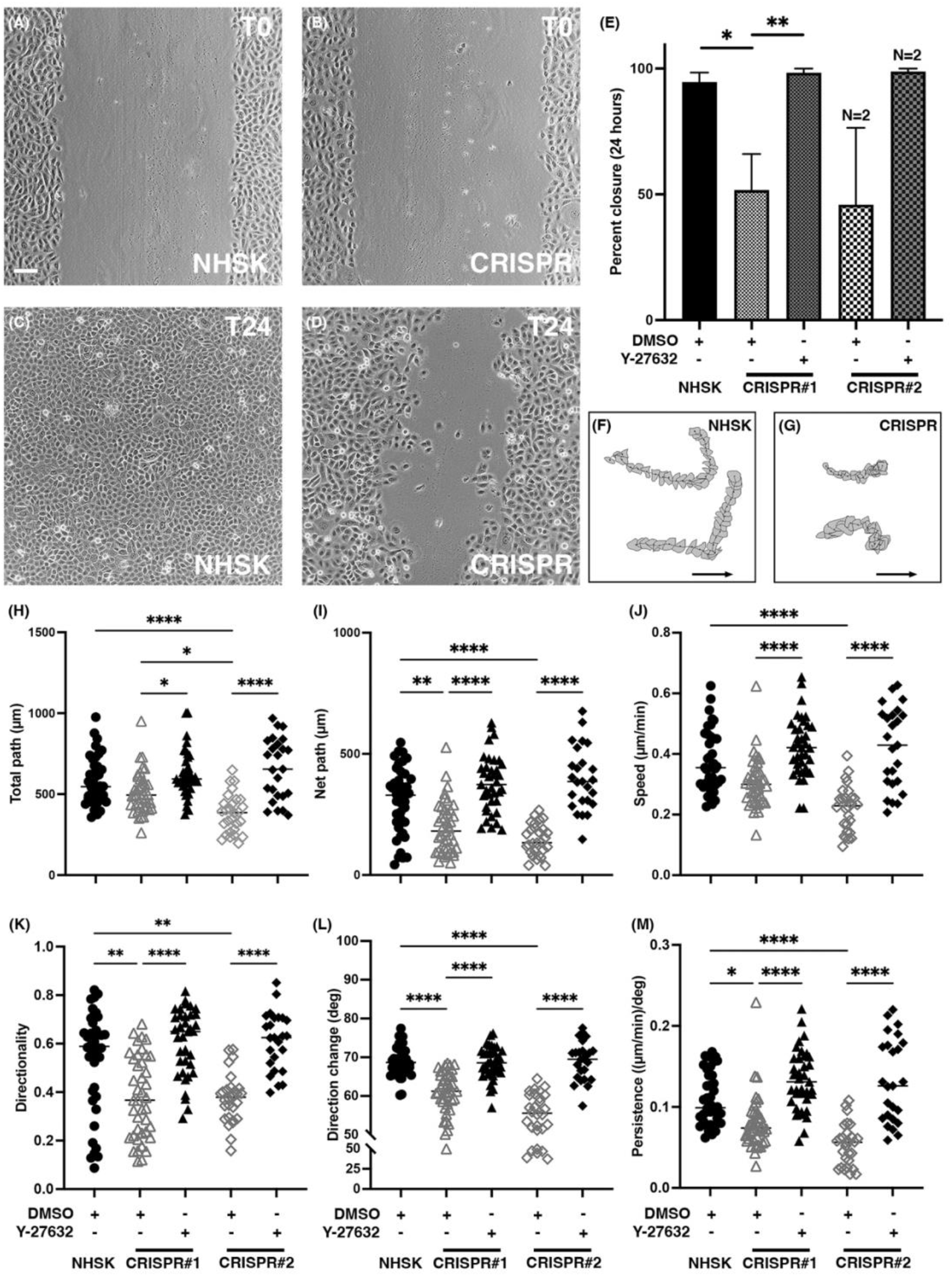
ARHGAP29 is required for single cell migration which is rescued by ROCK inhibition. A-D, Representative phase contrast micrographs of *in vitro* scratch wounds in confluent monolayers of NHSK (A,C) and CRISPR ARHGAP29 keratinocytes (B,D) grown in KSFM. Scale bar = 100 μm. T0 = 0 hours after scratch; T24 = 24 hours after scratch. E, Quantification of the percentage of scratch closure at 24 hours in CRISPR keratinocytes with or without Y-27632 treatment. Values are the means ± SEM, * *P* < 0.05 and ** *P* < 0.01 following ordinary one-way ANOVA with Tukey’s multiple comparisons *post-hoc* test. N = 3 per group unless otherwise noted. F and G, DIAS8-generated centroid tracks and stacked perimeter plots of representative NHSK (F) and CRISPR (G) keratinocytes where the arrow represents the direction of migration. H-M, Analysis of video recording and tracing of single keratinocytes during *in vitro* scratch wound closure with or without Y-27632. * *P* < 0.05, ** *P* < 0.01 and **** *P* < 0.0001 following Kruskal-Wallis test with Dunn’s multiple comparisons *post-hoc* test (only relevant comparisons are shown). N = 25-40 cells per group.

To assess which aspects of cellular migration are affected by the reduction of ARHGAP29, we performed single cell tracking analysis on time-lapse microscopy videos of *in vitro* scratch assays over the course of 24 hours. During that time, control keratinocytes showed processive single cell migration with few changes in direction (Figure 4F,H-M). However, CRISPR ARHGAP29 keratinocytes exhibited a significantly shorter net path, reduced directionality with more frequent direction changes and therefore reduced persistence (Figure 4G,H-M). Additionally, CRISPR#2 keratinocytes displayed a significant decrease in total path (Figure 4H) and speed (Figure 4J). Protein levels of ARHGAP29 in CRISPR#1 keratinocytes were higher than in CRISPR#2 keratinocytes, suggesting a potential dosage effect of ARHGAP29 on the severity of migration defects. Collectively, these results demonstrate that ARHGAP29 promotes path length, directionality and persistence of single cell keratinocyte migration during scratch closure *in vitro*.

### Inhibition of ROCK rescues migration defects in CRISPR ARHGAP29 keratinocytes

ARHGAP29 has previously been reported to act as a GAP with a high specificity for RhoA, but ARHGAP29 also interacts with Cdc42 and Rac1 *in vitro.^10,30^* To confirm that the defects observed in keratinocyte migration are due to alteration of the RhoA pathway, we treated control and CRISPR ARHGAP29 keratinocyte cell lines with Y-27632, an inhibitor of the RhoA effector protein ROCK.^38^ As described above, control keratinocytes grown in control conditions closed the scratch within 24 hours, while CRISPR ARHGAP29 keratinocytes showed a delay in scratch closure (Figure 4A-E). However, all cell lines closed the scratch within 24 hours in the presence of the ROCK inhibitor regardless of ARHGAP29 levels (Figure 4E).

To assess whether ROCK inhibition would also rescue other aspects of single cell migration, we performed single cell tracking analysis on DMSO treated (control, as described above) and Y-27632 treated keratinocytes throughout scratch closure. Indeed, ROCK inhibition rescued all measured migration defects in the CRISPR ARHGAP29 keratinocytes including total path length, net path length, speed, directionality, direction change and persistence (Figure 4H-M). These data suggest that the cell migration defects observed in the CRISPR ARHGAP29 keratinocyte cell lines are due to alteration of the RhoA pathway.

### ARHGAP29 is required for collective cell migration in vitro

Keratinocytes can migrate individually (single cell) or collectively, as they would *in vivo*. In contrast to single cell migration where a single cell moves freely (i.e. not attached to another cell) in a dish or tissue, collective cell migration involves a group of cells attached to each other by cell-cell adhesions and move together as a sheet.^39^ To further investigate the single cell migration defect observed in ARHGAP29 knockdown keratinocytes, we cultured keratinocytes using the classic method developed by Rheinwald and Green using an irradiated NIH3T3 feeder layer and DMEM:HAM-based medium.^40^ As these culture conditions favor proliferation, we treated keratinocytes with Mitomycin C 5 hours prior to administering the scratch and throughout subsequent closure. Similar to single cell conditions, we observed complete closure of the scratch in control cells after 12 hours (Figure 5A,D,G,H) and observed significant delays in scratch closure in ARHGAP29 knockdown cell lines (Figure 5B,E,G,H) despite no significant difference in original scratch area (data not shown). Further, the migration delay was rescued when ARHGAP29 was added back to sh#3 keratinocytes (Figure 5C,F,H). Together, these data demonstrate that ARHGAP29 is required for keratinocyte collective migration.

**Figure 5:**
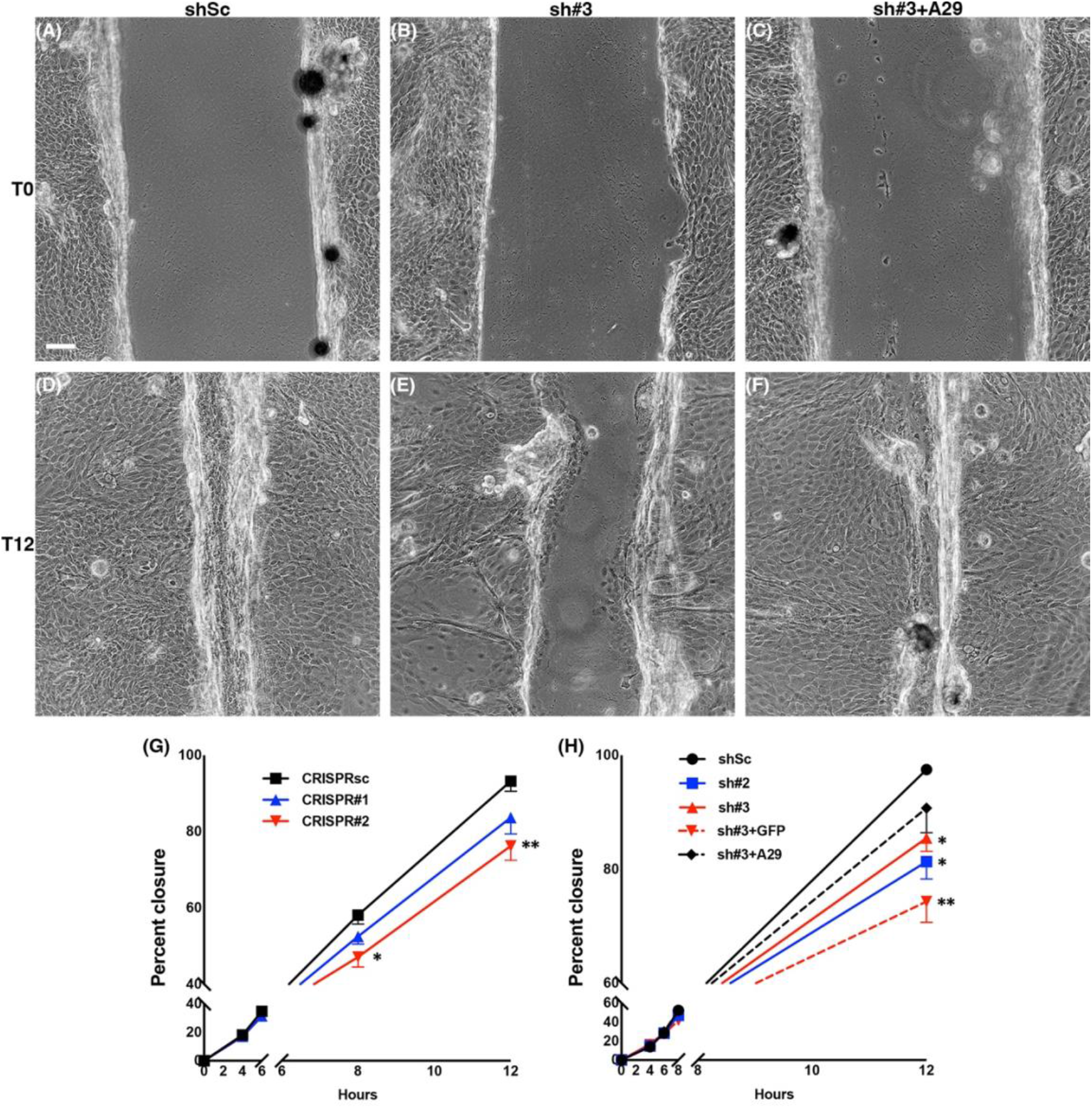
ARHGAP29 is required for collective cell migration. A-F, Phase contrast micrographs of *in vitro* scratch wounds in confluent monolayers of shRNA scramble (shSc; A,D), shRNA ARHGAP29 (sh#3; B,E), and sh#3 transduced with ARHGAP29 (sh#3+A29; C,F) keratinocytes grown in DMEM:HAM. Scale bar = 100 μm. T0 = 0 hours after scratch and T12 = 12 hours after scratch. G and H, Quantifications of the percentage of scratch closure over a 12 hour period in all CRISPR (G) and shRNA cell lines (H). Values are the means ± SEM, * *P* < 0.05 and ** *P* < 0.01 after two-way ANOVA with Tukey’s multiple comparisons *post-hoc* test. N = 6 per group.

## Discussion

In this study, we investigated the role of ARHGAP29 in keratinocyte morphology, proliferation and migration. Using novel keratinocyte cell lines knocked down for ARHGAP29, we demonstrate that these cells exhibit increased filamentous actin (stress fibers), phospho-myosin regulatory light chain (contractility), larger cell area and population doubling time (proliferation). Furthermore, we report that ARHGAP29 knockdown keratinocytes displayed impaired single cell and collective cell migration, and that this phenotype is specific to ARHGAP29 and the RhoA pathway.

*ARHGAP29* has been knocked down using an RNAi approach in fibroblasts, endothelial, and cancer cells^15,17,37,41^, and a CRISPR/Cas9 strategy in HEK293A cells.^42^ Our studies demonstrate that ARHGAP29 levels can also be reduced in keratinocytes. However, we were unable to completely knockout ARHGAP29 in these cells, leaving us to speculate that this enzyme may be required for keratinocyte survival. In fact, one of our shRNA cell lines (sh#1) never expanded in culture. Compared to the other cell lines in which ARHGAP29 was knocked down, our keratinocytes are minimally transformed and overall closer to their original parental line. As such, their requirement for survival may be different from fully transformed cells, including a requirement for ARHGAP29. Our CRISPR/Cas9 approach led to mutations in *ARHGAP29* predicted to cause premature truncation mutations within the first coding exon. In theory, this should have resulted in a total loss of ARHGAP29 protein due to nonsense mediated decay of the mutated transcript, yet we continued to detect some levels of ARHGAP29 by Western blot. Similar phenomena have been reported throughout the literature in association with the CRISPR/Cas9 system, in which all alleles of a gene are modified, but the protein is still expressed.^43,44^ Given that the antibodies we used were generated using an epitope at the C-terminus of the ARHGAP29 protein and the bands observed in controls and *ARHGAP29* knockdown lines were all near the expected size of ARHGAP29 (142 kDa), we speculate that keratinocytes may adopt an alternative translation start site downstream of the CRISPR edits resulting in an in-frame ARHGAP29 protein. Use of this alternate start site would likely result in a protein lacking a small portion of the N-terminus and still maintain some, if not all, degree of function. Testing this hypothesis would involve performing mass spectrometry on purified ARHGAP29 protein from the ARHGAP29 CRISPR cells as previously suggested.^43^

The most striking phenotype of ARHGAP29 knockdown keratinocytes was their larger cell area and increase in population doubling time. Given that ARHGAP29 negatively regulates RhoA and active RhoA promotes actin polymerization,^5^ it was not surprising that ARHGAP29 knockdown keratinocytes exhibited increased filamentous actin (stress fibers) and pMLC. Despite being consistent with an increase in RhoA activity, these observations differ from the phenotype of IRF6-deficient keratinocytes, in that reduction of IRF6 increased their proliferative capacity despite their large size and increased filamentous actin. IRF6 is a transcription factor with many targets critical for keratinocyte differentiation. Therefore, loss has far-reaching effects on cell proliferation. Current data does not support *ARHGAP29* as a direct transcriptional target for IRF6^45^, further supporting that ARHGAP29 mediates only some IRF6 effects, however which ones and how would remain to be formally tested. Interestingly, one IRF6 target is Grainyhead-like 3 (*GRHL3*), another pro-differentiation transcription factor, which regulates RhoGEF19, a GEF which activates RhoA.^45–47^ Similar to IRF6 and ARHGAP29, knockdown of *GRHL3* in keratinocytes lead to delayed scratch closure and impaired cell migration.^47^ Since ARHGAP29 and RhoGEF19 display opposing functions in RhoA cycling yet show the same phenotype, it highlights the importance of its cyclical regulation in migration.

Throughout our studies, we noticed an apparent correlation between the severity of defects (e.g., migration, cell size) and the amount of ARHGAP29 knockdown. Consistent with this observation, heterozygosity of a missense genetic variant in *ARHGAP29* identified in a multiplex family with cleft palate lead to the amino acid change p.Ser552Pro and reduced ARHGAP29 expression. Interestingly, the corresponding mutated protein failed to promote keratinocyte migration compared to wild-type ARHGAP29.^22,48^ Dosage effect of ARHGAP29 is also evident *in vivo*, as mice heterozygous for *Arhgap29* exhibit transient oral adhesions during palatogenesis, a characteristic recognized in the field as a risk factor for cleft lip with or without palate, while homozygous *Arhgap29* null animals die early during embryogenesis.^12,49^ Interestingly, while decreased ARHGAP29 levels have been associated with cleft lip with or without palate, increased ARHGAP29 levels are also correlated with poor prognosis in a number of cancers including gastric, breast, prostate and liver due to increased proliferation, migration, invasion and metastasis.^16–20^ Collectively, these studies support a model in which too much or too little ARHGAP29 has detrimental effects on proliferation and migration in the context of human development and disease.

In summary, our study describes a novel role for ARHGAP29 in keratinocyte proliferation and migration, two processes also required for tissue repair. It is worth noting that *ARHGAP29* transcript was upregulated 6 hours following an *in vitro* scratch wound of keratinocytes and was classified as “signal transducer,” suggesting its potential role in inhibiting RhoA activity during tissue repair.^50^ ARHGAP29 level was also increased in myofibroblasts under hypoxic conditions,^41^ another characteristics of wounds. Further studies evaluating the specific role of ARHGAP29 in wound healing may provide insights into its broader effect in tissue repair. As a number of RhoA/ROCK pathway inhibitors are currently being tested in clinical trials, including for the treatment of spinal cord injury,^51^ it would be interesting to determine if ARHGAP29 contributes to the effects of these drugs for future therapies.

## Experimental Procedures

### Keratinocyte culture

TERT-Cherry N-HSK-1 (Normal Human Skin Keratinocytes immortalized with a TERT-Cherry overexpression, a generous gift from Dr. Klingelhutz^29^) henceforth referred to as NHSKs were grown in Keratinocyte-SFM (1X) media (Gibco Thermo Fisher Scientific, Waltham, MA) with 0.16 μg/L EGF, 25 μg/mL BPE, and 1% Penn-Strep (Gibco Thermo Fisher Scientific) with the exception of the colony forming efficiency (CFE) and collective cell *in vitro* scratch assays in which keratinocytes were grown on an irradiated NIH3T3 feeder layer following the method of Rheinwald and Green as previously described.^29^ Cells were maintained in a tissue culture incubator at 37°C and 5% CO_2_. Population doubling time (PDT) was calculated using the following formula: PDT = (H/(3.32(log A – log B)) where H = hours between seeding cells and passaging, A = the final number of cells counted at passaging and B = the number of cells seeded.

### CRISPR Lentiviral transduction

Benchling.com was used to design guide RNAs targeting exon 2 of *ARHGAP29*. The ARHGAP29 CRISPR#1 lentivirus was generated by the University of Iowa Viral Vector Core using a LentiCRISPR_v2 plasmid modified to contain a gRNA targeting exon 2 of *ARHGAP29* 5’caccGATATTACAACTTCTGAAATG3’.^52^ The ARHGAP29 CRISPR#2 and scrambled (CRISPRsc) lentivirus were generated in our laboratory by transducing the following plasmids into TSA cells: pPAX2, pVSVG, and a LentiCRISPR_V2 plasmid modified to include either the gRNA targeting exon 2 of *ARHGAP29* 5’caccgATCCGGATCAAAAATAGAGT’ or the scrambled gRNA 5’GGGCCCGCATAGGATATCGC3’. For all viruses, NHSKs were treated for six hours with Polybrene (10 μg/ml) and viral particles before changing media back to Keratinocyte-SFM. Two days following transduction, puromycin selection was performed to remove any colonies not expressing the lentiviral construct. To generate monoclonal lines, keratinocytes were plated at very low density into 100 mm dishes and allowed to expand until small colonies were formed. Ring cloning was then performed by isolating and passaging individual colonies into separate dishes for expansion.

### TOPO cloning and sequencing

The second exon of ARHGAP29 was sequenced using the following primers: forward: 5’TGGCATCAAGGCAGATACAG3’, reverse: 5’ACCCAACCACAGGAATCAAA3’. Due to the majority of the cell lines being compound heterozygotes, the alleles had to be isolated using TOPO cloning. The PCR product of the primers listed above was ligated into the pCR™4-TOPO^®^ vector (Invitrogen), and transformed into One Shot^®^ TOP10 *E. Coli*. Colonies (6-12) from each cell line were selected, expanded, and plasmids were isolated using a High-Speed Plasmid Mini Kit (IBI Scientific, Dubuque, IA). Plasmids were then Sanger sequenced using the primers listed above. Sequence analysis was performed using Benchling.com software.

### shRNA Lentiviral transduction

VectorBuilder’s shRNA Target Design tool was used to design three different short hairpin RNAs with high knockdown scores for *ARHGAP29*. The lentiviral vectors used to knockdown *ARHGAP29* in our study were constructed and packaged by VectorBuilder. The following vector names and IDs may be used to retrieve detailed information about the vector on vectorbuilder.com: pLV[shRNA]-Puro-U6>hARHGAP29[shRNA#1] (Vector ID: VB200721-1456nau) (sh#1), pLV[shRNA]-Puro-U6>hARHGAP29[shRNA#2] (Vector ID: VB200721-1457ghg) (sh#2), pLV[shRNA]-Puro-U6>hARHGAP29[shRNA#3] (Vector ID: VB200721-1458eng) (sh#3), pLV[shRNA]-Puro-U6>Scramble_shRNA#1 (Vector ID: VB010000-0005mme) (shSc). For all viruses, NHSKs were treated overnight with Polybrene (4 μg/ml) and viral particles (multiplicity of infection ~15). The following day transduced cells were passaged and re-plated in Keratinocyte-SFM with puromycin (1 μg/ml) for 7 days.

### Lentiviral transduction of ARHGAP29 (“rescue”)

The “ARHGAP29 rescue” lentivirus was generated by the University of Iowa Viral Vector Core using a pFIV3.2CMVmcswtIRESzeocin^R^ plasmid modified to contain an N-terminal GFP tag on the human *ARHGAP29* cDNA. Similar plasmid only containing eGFP was used as a control. NHSK cells modified with sh#3, which targets the 3’ untranslated region of the human *ARHGAP29* cDNA, were treated with Polybrene (4 μg/ml) and either the ARHGAP29-GFP (rescue) or eGFP (control) lentiviruses overnight before being replaced with Keratinocyte-SFM the following day. Cells were allowed to expand for 48 hours post-transduction before bulk sorting GFP positive populations with a FACSAria Fusion flow cytometer (Becton Dickinson).

### Western blot analysis

Keratinocytes near confluency were extracted using 2X Laemmli Buffer.^53^ Samples were then loaded on a SurePAGE 4-12% Bis-Tris SDS-PAGE gel (Genscript, Piscataway, NJ) under denaturating conditions followed by protein transfer onto Immuno-Blot^®^ PVDF Membranes (Bio-Rad Laboratories, Hercules, CA). Membranes were blocked in 10% non-fat dry milk for two hours at room temperature (Hy-Vee, Des Moines, IA), and incubated in primary antibody diluted in Tris-buffer salt with 0.1% Triton X-100 overnight at 4°C under rocking conditions. Membranes were then incubated with horseradish peroxidase-conjugated secondary IgG antibodies. Immunoblot visualization was performed with the Amersham™ Imager 600 (GE Healthcare, Uppsala, Sweden) following chemiluminescent detection (SuperSignal^®^ West Pico PLUS and West Femto Chemiluminescent Substrate (Thermo Scientific, Rockford, IL).

### Immunofluorescence microscopy

Keratinocytes were plated on glass coverslips and grown in Keratinocyte-SFM until they reached 30-50% confluency. Cells were fixed in 4% paraformaldehyde for 10 minutes and permeabilized using 0.2% Triton X-100 in PBS for 2 minutes. Fixed cells on coverslips were blocked using 3% Normal Goat Serum (Vector Laboratories, Burlingame, CA) in 2% PBS-Bovine Serum Albumin for 30 minutes at room temperature. Following blocking, cells were incubated in primary antibody for one hour at room temperature. The coverslips were then washed with 1X PBS and incubated with secondary antibody for one hour at room temperature. Coverslips were mounted with ProLong^TM^ Diamond Antifade Mountant (Thermo Fisher Scientific, Waltham, MA). Images were collected with a Zeiss 880 Confocal Microscope using ZEN Software (Zeiss, Oberkochen, Germany). Confocal images were processed using Fiji software.^54^

### Antibodies

The following antibodies were used for immunofluorescence: Phalloidin-TRITC (Sigma-Aldrich, St-Louis, MO; catalog # P1951) was used at 1/10,000; mouse monoclonal against Vinculin (Sigma-Aldrich; catalog # V9264) used at 1/300; rabbit polyclonal against Phospho-Myosin Regulatory Light Chain 2 at position Ser19 (Cell Signaling, Danvers, MA; catalog # 3671) was used at 1/100; Alexa Fluor 488 goat anti-mouse and anti-rabbit IgG (Invitrogen, Waltham, MA; catalog # A-11001 (mouse) and A-11008 (rabbit)) were used at 1/200. Hoechst (bisBenzimide H 33342 trihydriochloride; Sigma-Aldrich) powder was dissolved at 1 mg/mL in ddH2O and used at 1/10,000.

The following antibodies were used for Western blotting: Rabbit polyclonal against ARHGAP29 (Novus Biologicals, Littleton, CO; catalog # NBP1-05989) was used at 1/1,000; mouse monoclonal against GAPDH (Ambion; Gibco Thermo Fisher Scientific; catalog # AM4300) was used at 1/9,000; mouse anti-rabbit IgG-HRP (eBiosciences (San Diego, CA; catalog # 18-8816-33) was used at 1/5,000; sheep anti-mouse IgG-HRP (GE Healthcare, Chicago, IL; catalog # NA931V) was used at 1/5,000.

### Live imaging, single cell in vitro scratch assay and migration analysis

Phase contrast micrographs of confluent keratinocytes were collected using an Olympus IX-81 inverted microscope (Olympus, Shinjuku, Tokyo, Japan) equipped with Slidebook 6 software (Intelligent Imaging Innovations, Denver, CO). Confluent keratinocytes were scratched as previously described and fed with either Keratinocyte-SFM + DMSO or Keratinocyte-SFM + 10 mM Y-27632 (Sigma).^6^ The open area of the scratch was analyzed in three different areas and averaged using Fiji at 0 h, 2 h, 4 h, 6 h, 8 h, 12 h, and 24 h following scratching.^54^ Scratches were performed on 3 independent passages per cell line.

Time lapse images collected from the IX-81 were compiled into videos and analyzed using the DIAS8 software as previously described.^6,55,56^ For each video, 10 cells were selected from within a 3 cell-depth of the scratch margin. Manual tracing of the perimeter of the cells throughout the video was performed. The data were analyzed using DIAS8 software, which allowed the quantification of the net path, total path, speed of migration, directionality, persistence, and roundness of individual keratinocytes.

### Collective cell migration

Keratinocytes were grown to confluency on an irradiated NIH3T3 feeder layer in a DMEM:HAM-based medium as previously described.^40^ Cells were treated with Mitomycin C (5 μg/mL) for 5 hours prior to performing a scratch as previously described.^6^ After scratching, medium was aspirated and replaced with fresh DMEM:HAM + 5 μg/mL Mitomycin C. Phase contrast micrographs were collected using a Nikon Eclipse Ts2 microscope equipped with a Nikon DS-Ri2 camera operated by the NIS-Elements Imaging Software (Minato, Tokyo, Japan). Scratches were photographed at 0 h, 4 h, 6 h, 8 h, 12 h, and 24 h following scratching. For each micrograph, 3 different locations of the open area were analyzed and the distance between scratch edges measured and averaged using Fiji.^54^ Six scratches from 3-4 different passages for each cell line were analyzed.

### Colony forming efficiency assays

Colony forming efficiency (CFE) was assessed 12 days following the plating of 500 keratinocytes per 60 mm dish as previously described.^29^ Three technical replicates were plated at each of 3 different passages for each cell line analyzed. Technical replicates were averaged for each cell line at each passage for a final N=3. On day 12, dishes were washed twice with PBS, stained with a 75% *v/v* Giemsa Stain (RICCA, Arlington, TX) in methanol solution, then washed with Giordano buffer (0.66% *w/v* potassium phosphate monobasic, 0.32% *w/v* sodium phosphate dibasic, 0.1% *v/v* formaldehyde 37% in ddH2O) followed by two final washes with tap water. Dishes were allowed to dry upside-down and photographed. Colonies in the dish were counted and their area quantified using NIH-ImageJ software.

## Acknowledgements

The authors would like to thank Drs. Eric Taylor, Adam Rauckhorst and Botond Bonfi for critical discussion related to CRISPR, and all Dunnwald lab and Tootle lab members for continuous input. The technical expertise of Annemarie Carver, Campbell Mitvalsky and members of the W.M. Keck Dynamic Image Analysis facility in the Biology department of the University of Iowa is greatly appreciated. The data presented herein were obtained using the Flow Cytometry Facility, University of Iowa Central Microscopy Research Facility, University of Iowa Viral Vector Core, and the Genomics Division of the Iowa Institute of Human Genetics which are all supported by the Carver College of Medicine at the University of Iowa. This work was supported by funding from the National Institute of Health (NIH, R01-AR067739, MD; T32-GM145441, EA) and the American Association for Anatomy Fellows Grant Program (MD).

---

**Supplemental Figure 1: ARHGAP29 CRISPR genome editing validation.** ARHGAP29 sequencing alignments generated using Benchling.com for NHSK, CRISPRsc, and the alleles of CRISPR#1 and CRISPR#2 as determined by TOPO cloning. A, DNA sequence where a dash (-) represents a single base pair deletion. B, Amino acid sequence where an asterick * represents a premature truncation mutation.

