## Supplementary figures and images for "ARHGAP29 is required for keratinocyte proliferation and migration"

### Supplemental Table 1

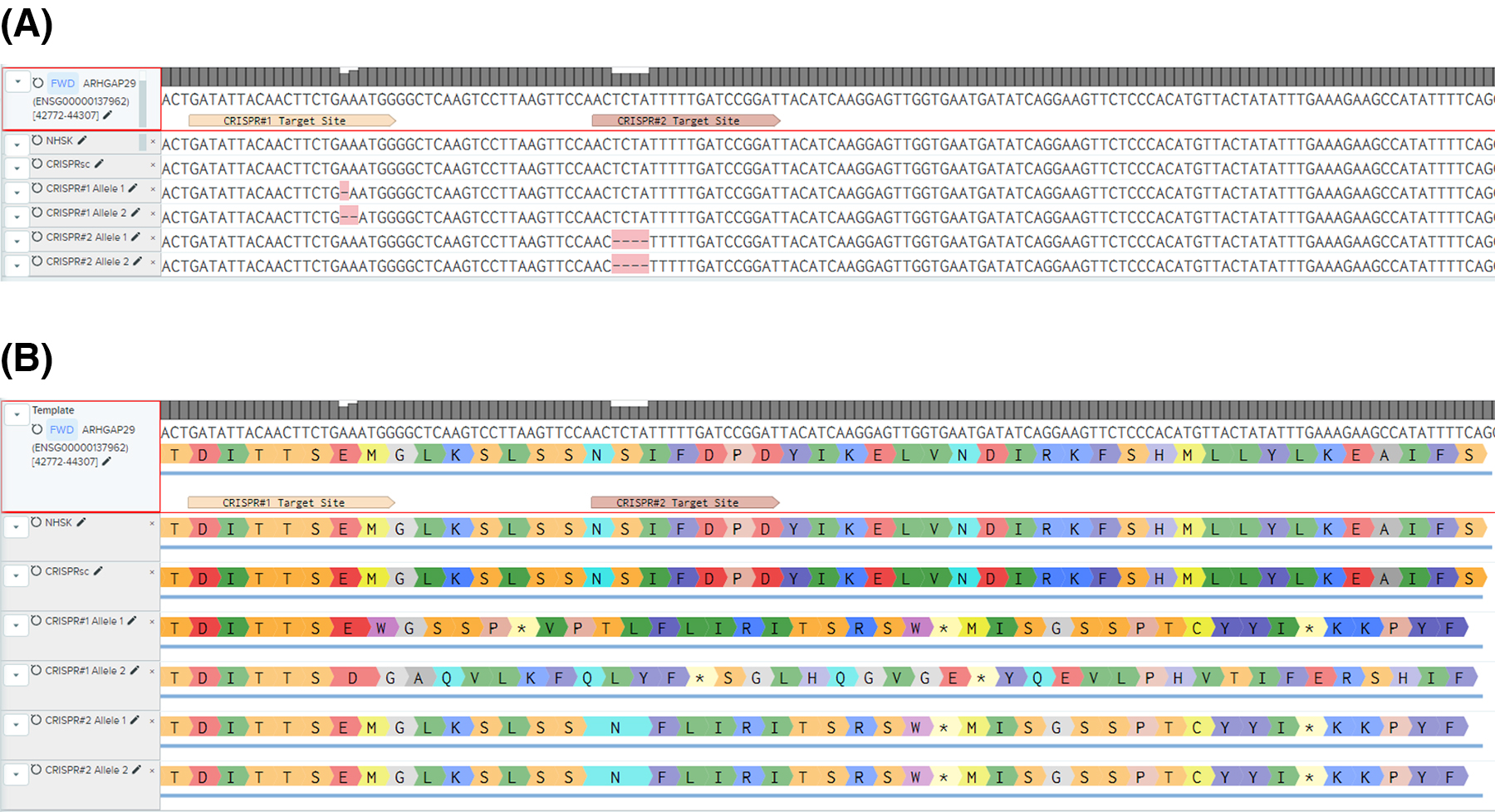
